# Network integration of multi-tumour omics data suggests novel targeting strategies

**DOI:** 10.1101/146209

**Authors:** Ítalo Faria do Valle, Giulia Menichetti, Giorgia Simonetti, Samantha Bruno, Isabella Zironi, Danielle Fernandes Durso, José C M Mombach, Giovanni Martinelli, Gastone Castellani, Daniel Remondini

## Abstract

We characterize different tumour types in the search for multi-tumour drug targets, in particular aiming for drug repurposing or novel drug combinations. Starting from 11 tumour types from The Cancer Genome Atlas, we obtain three clusters based on transcriptomic correlation profiles. A network-based analysis, integrating gene expression profiles and protein interactions of cancer-related genes, allowed us to define three cluster-specific signatures, with genes belonging to NF-ĸB signaling, chromosomal instability, ubiquitin-proteasome system, DNA metabolism, and apoptosis biological processes. These signatures have been characterized by different approaches based on mutational, pharmacological and clinical evidences, demonstrating the validity of our selection. Moreover, we defined new pharmacological strategies validated by *in vitro* experiments that showed inhibition of cell growth in two tumour cell lines, with significant synergistic effect. Our study thus provides a list of genes and pathways with the potential to be used, singularly or in combination, for the design of novel treatment strategies.

## Introduction

High-throughput molecular profiling has changed the approach to study cancer. For decades, anatomical localization and histological features have guided the identification of cancer subtypes, but the genomic profiling of tumour samples has revealed differences and similarities that go beyond the histopathological classification. The diversity in genomic alteration patterns often stratifies tumours from the same organ or tissue, while tumours in different tissues may present similar patterns^1–3^. For example, mutational profiling of transcription factors/regulators show tissue specificity, while histone modifiers can be mutated similarly across several cancer types^4^. Hoadley et. al.^2^ suggests that lung squamous, head and neck, and a subset of bladder cancers form a unique cancer category typified by specific alterations, while copy number, protein expression, somatic mutations and activated pathways divide bladder cancer into different subtypes. The analysis of cancer transcriptomes revealed that the same tumour may originate from several cell types, and different biological processes may lead to malignant transformation^4^. Moreover, similar pathways may be activated in different cancers, like ovarian, endometrial and basal-like breast carcinomas^6,7^.

Notwithstanding the enormous increase of knowledge on tumour processes, actually, a practical application of this knowledge to new treatment strategies has not advanced with the same pace. For example, common genetic alterations can predict similar responses to pharmacological therapies across multiple cancer cell lines^8–10^, thus such common molecular and functional profiles could enable the repurposing of therapies from one cancer to another.

The huge amount of heterogeneous types of data for a large number of tumours requires novel approaches capable to integrate such information into a unified framework: for this aim, we propose a study of gene networks based on expression profiling and mutational data, in combination with cancer-specific functional annotation. Starting from whole-genome transcriptional profiling extracted from The Cancer Genome Atlas (TCGA) data portal (https://gdc-portal.nci.nih.gov/), we selected a curated subset of cancer-related genes and pathways described in the Ontocancro database (http://ontocancro.inf.ufsm.br/), and mapped these data onto the BioPlex protein-protein interaction network^11^. A structural analysis of the obtained networks, based on node centrality, allowed us to rank their relevance and to obtain specific signatures, that may provide multi-tumour drug targets, prognostic markers, and a molecular taxonomy for effective cancer categorization.

The validation of our signatures through literature interrogation, clinical information and by *in vitro* testing, makes us confident that this study can help both clinical and research communities, providing novel targets for multi-drug approaches and for repurposing of existing drugs.

## Results

We analyzed transcriptomic data of 2378 samples from 11 tumour types (Table 1) considering 760 cancer-related genes with protein-protein interaction annotation (Bioplex-Ontocancro network, see Methods). The tumour datasets were clustered in three groups based on their gene-gene correlation matrices (see Methods) containing, respectively, 2, 6 and 3 cancer types: 1) Colon adenocarcinoma (COAD) and Rectum Adenocarcinoma (READ); 2) Lung Adenocarcinoma (LUAD), Lung Squamous Cell Carcinoma (LUSC), Glioblastoma Multiforme (GBM), Ovarian Serous Cystadenocarcinoma (OV), Breast Invasive Carcinoma (BRCA), and Uterine Corpus Endometrial Carcinoma (UCEC); and 3) Brain Lower Grade Glioma (LGG), Kidney Renal Clear Cell Carcinoma (KIRC), and Kidney Renal Papillary Cell Carcinoma (KIRP) (Figure 1). By superimposing the correlation matrices (specific to each cluster) onto the BioPlex-Ontocancro network (common to all tumours), we obtained three weighted networks with approximately 80% of the original nodes and 60% of the original edges (Table 2, see Supplementary Figures 1-4).

**Figure 1.**
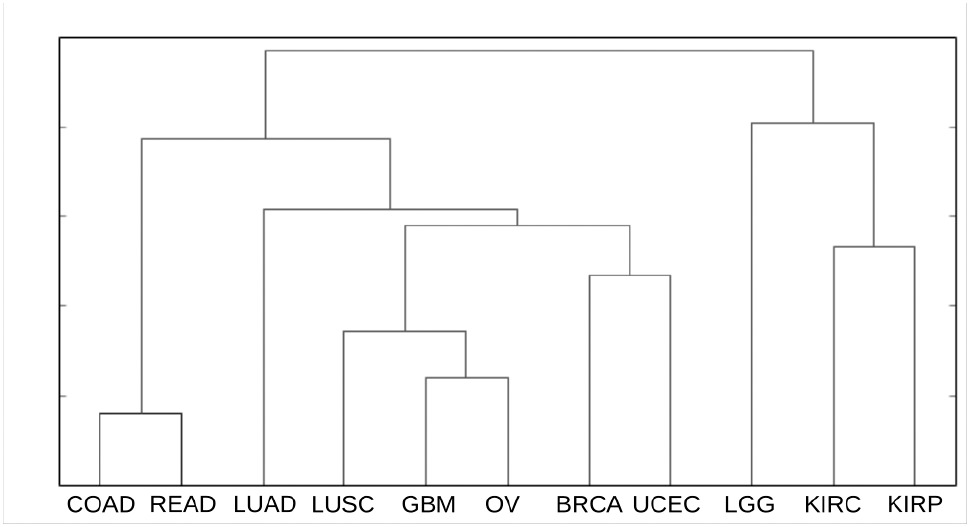
tumour clustering. For each tumour, we produced a matrix from the correlation (Pearson) of the expression profiles among 760 genes. The correlations values were adjusted by the CLR algorithm. Then, we clustered the resulting matrices by euclidean metrics.

**Table 1.** The datasets. List of tumours and their respective number of gene expression arrays

| Abbreviation | Cancer | Number of patients |
| --- | --- | --- |
| BRCA | Breast invasive carcinoma | 593 |
| COAD | Colon adenocarcinoma | 172 |
| GBM | Glioblastoma multiforme | 595 |
| KIRC | Kidney renal clear cell carcinoma | 72 |
| KIRP | Kidney renal papillary carcinoma | 16 |
| LGG | Brain lower grade glioma | 27 |
| LUAD | Lung adenocarcinoma | 32 |
| LUSC | Lung squamous cell carcinoma | 155 |
| OV | Ovarian serous cystadenocarcinoma | 590 |
| READ | Rectum adenocarcinoma | 72 |
| UCEC | Uterine corpus endometrial carcinoma | 54 |
| Total |  | 2378 |

**Table 2.**
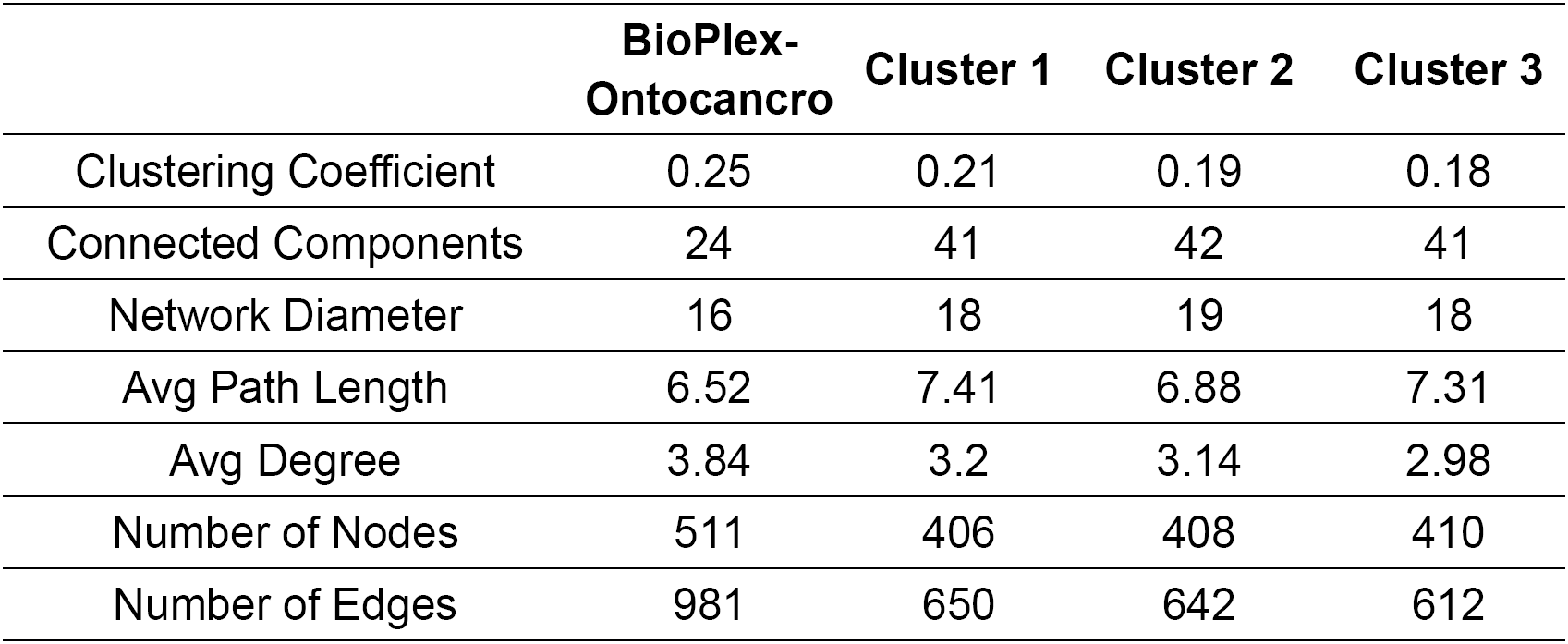
Network Properties. The table shows the main topological features of the cluster networks. Cluster 1: COAD and READ; Cluster 2: LUAD, LUSC, GBM, OV, BRCA, and UCEC; and Cluster 3: LGG, KIRC and KIRP.

|  | BioPlex-Ontocancro | Cluster 1 | Cluster 2 | Cluster 3 |
| --- | --- | --- | --- | --- |
| Clustering Coefficient | 0.25 | 0.21 | 0.19 | 0.18 |
| Connected Components | 24 | 41 | 42 | 41 |
| Network Diameter | 16 | 18 | 19 | 18 |
| Avg Path Length | 6.52 | 7.41 | 6.88 | 7.31 |
| Avg Degree | 3.84 | 3.2 | 3.14 | 2.98 |
| Number of Nodes | 511 | 406 | 408 | 410 |
| Number of Edges | 981 | 650 | 642 | 612 |

We hypothesized that the most central genes in each network should play a fundamental role in the tumours represented in the cluster. To find the most central genes we measured the Spectral Centrality (SC)^12^, related to the changes in network global diffusivity by node perturbation through a Laplacian formalism, and considered the nodes with SC above the 90^th^ percentile (25, 27 and 24 genes for clusters 1,2, 3 respectively, Table 3). We remark that the chosen signatures have only a small overlap with the most central nodes on the original “full” Bioplex-Ontocancro network not filtered by the cluster-specific correlation matrices (3/25, 13/27 and 4/24 common genes for clusters 1,2, 3, respectively) showing how the information on gene expression profile is highly specific for the considered tumour clusters. The top-ranking nodes also differ significantly from those obtained from other centrality measures such as degree and betweenness centrality (see Supplementary table 5). Moreover, even if some signature genes overlap between clusters, their links are different (Figures 2, 3, 4, and Supplementary Figure 5) evidencing a specific interaction pattern.

**Figure 2.**
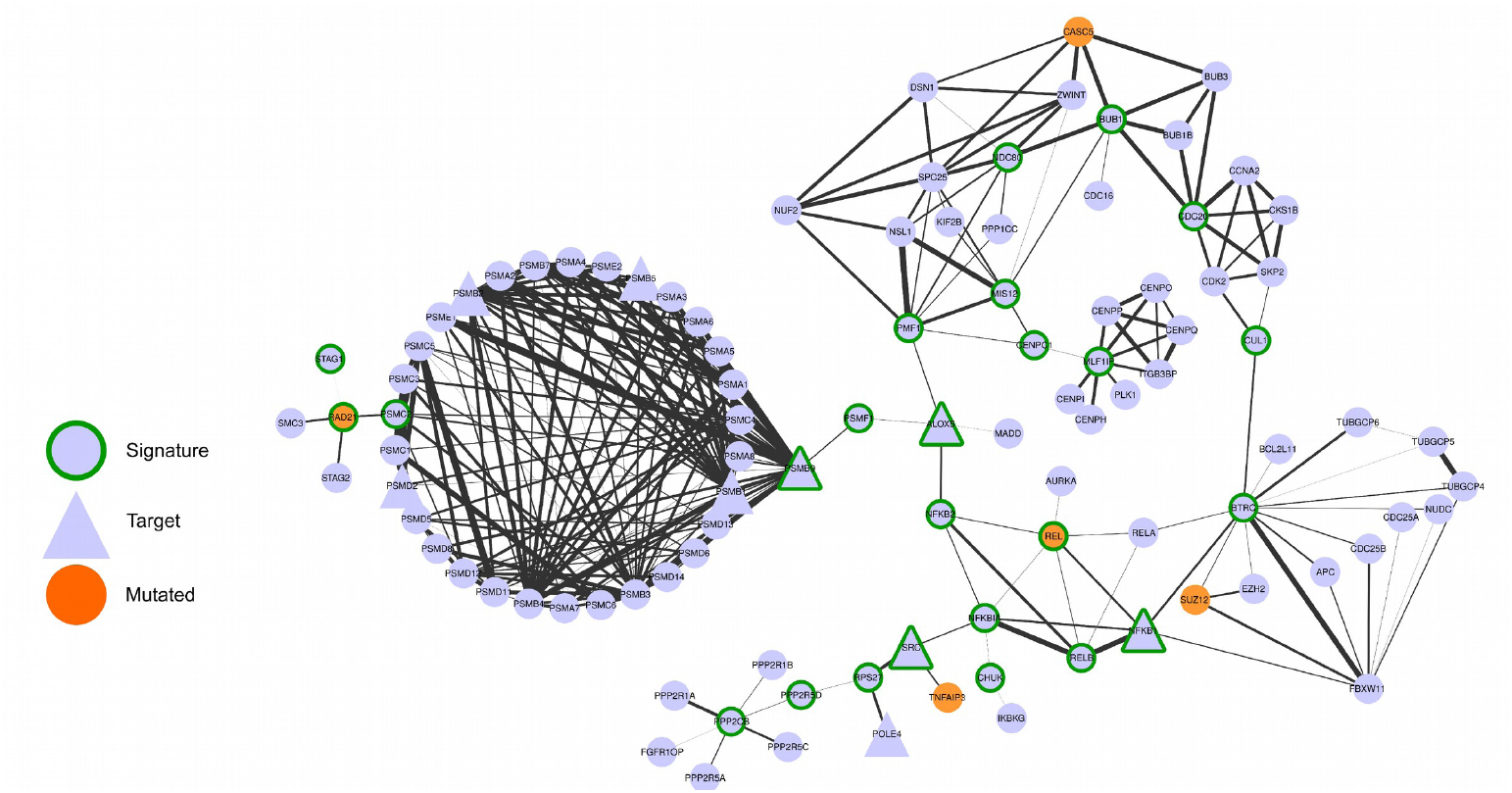
– Network composed by the first neighbors of the cluster 1 signature genes

**Figure 3.**
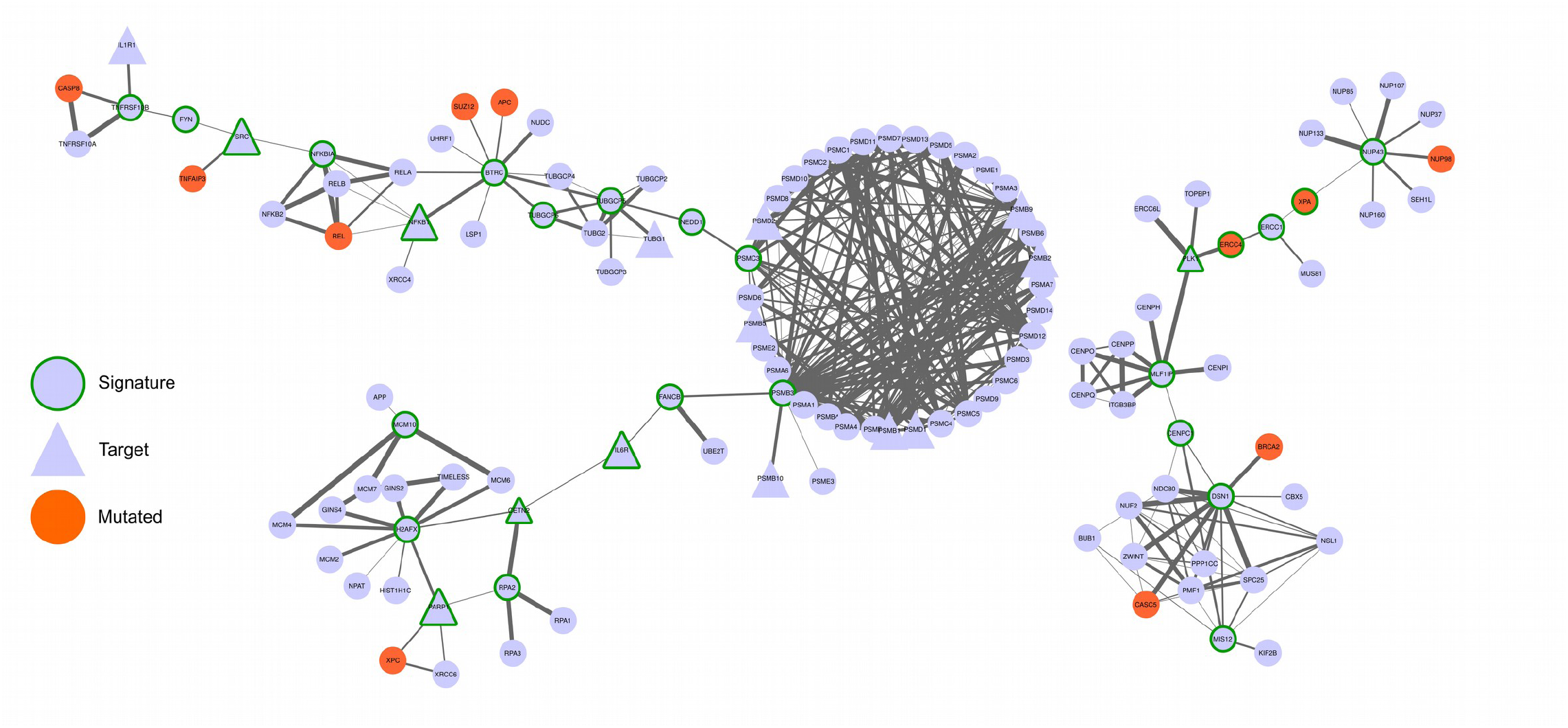
– Network composed by the first neighbors of the cluster 2 signature genes.

**Figure 4.**
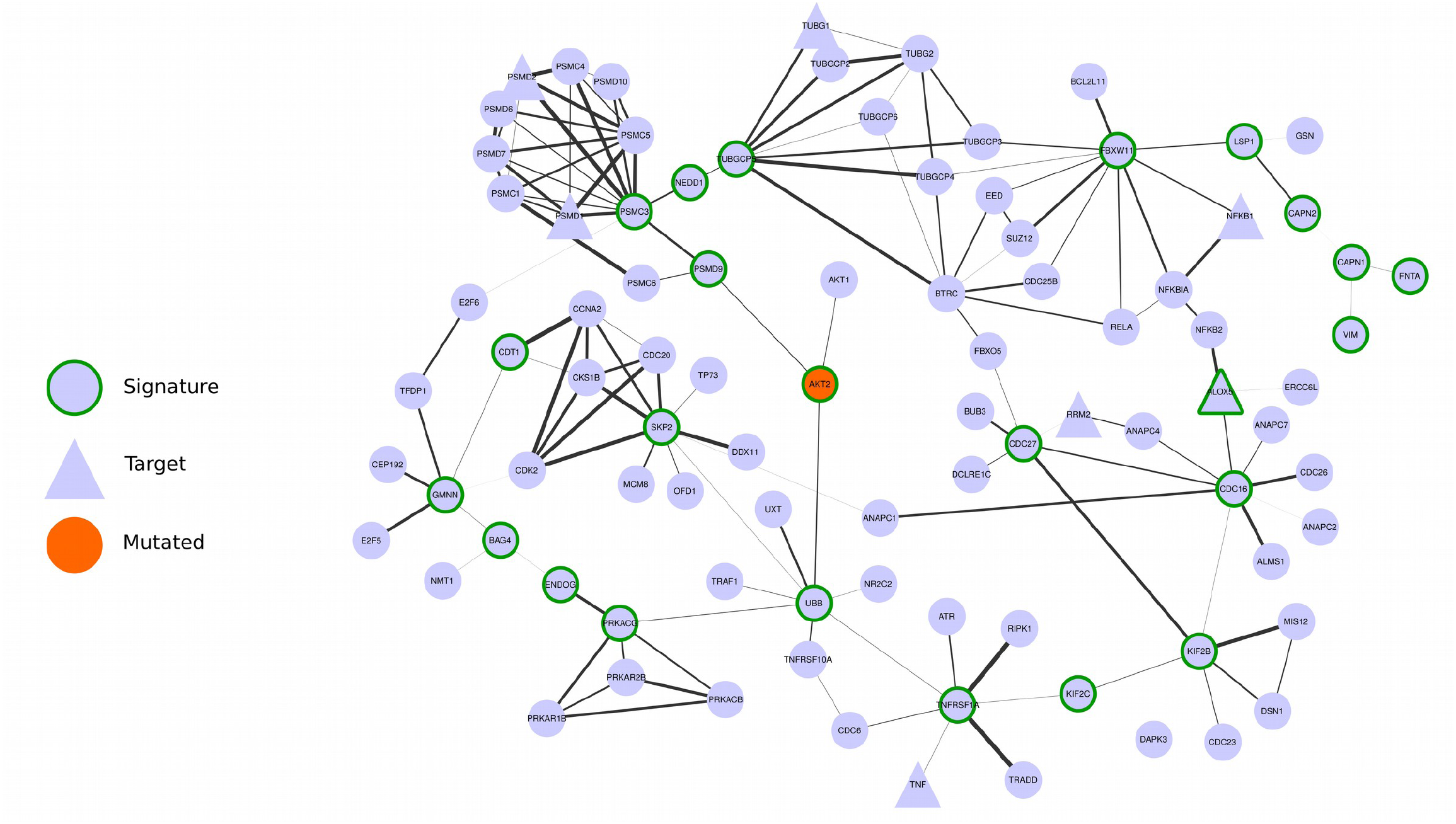
– Network composed by the first first neighbors of the cluster 3 signature genes

**Table 3.**
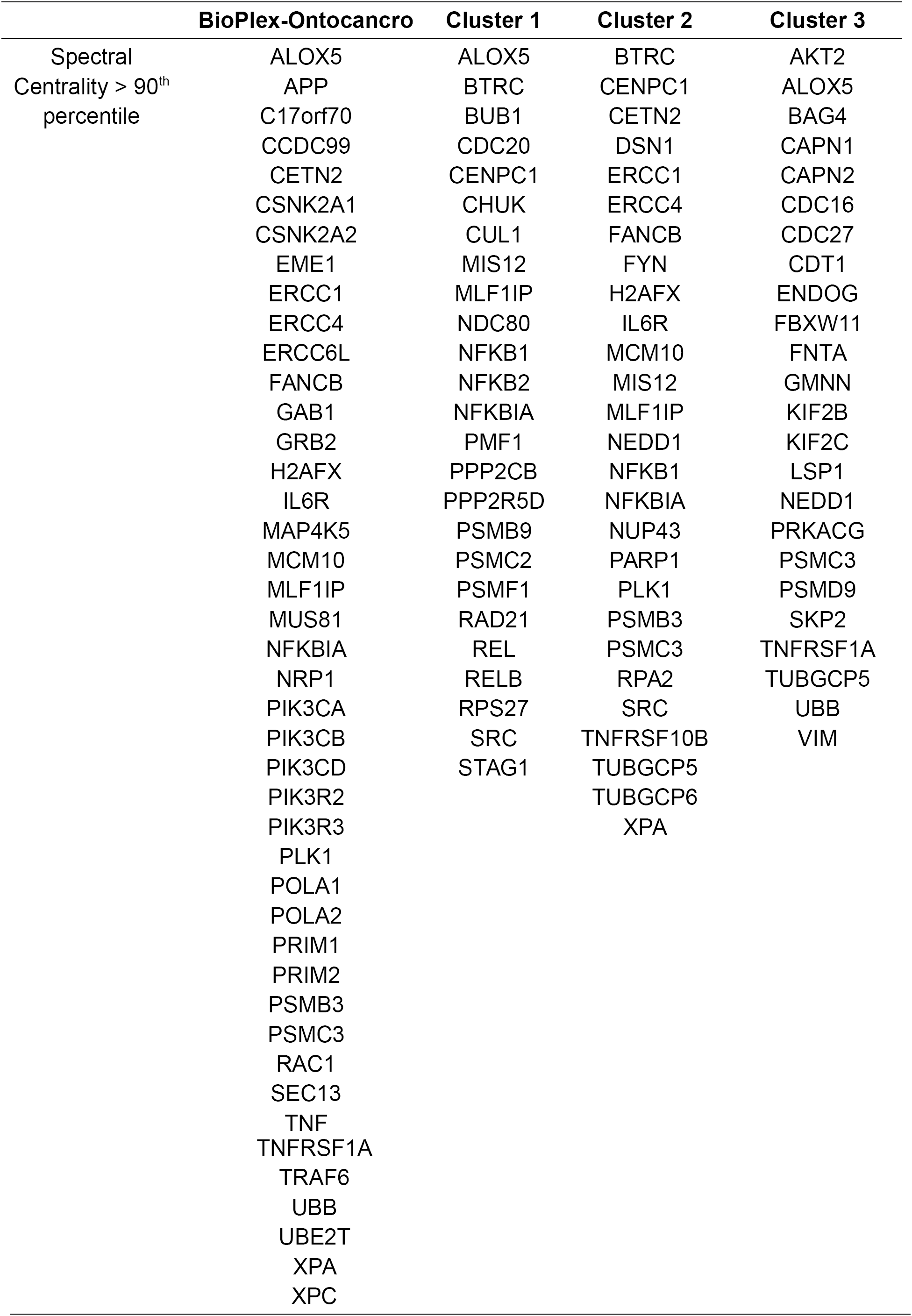
Gene Signatures. List of signature genes for the three tumour clusters.

We observed that all signatures contain genes related to three biological categories: NF-ĸB signaling pathway, chromosomal instability and ubiquitin-proteasome system (Table 4). The chromosomal instability category relates to genes involved in kinetochore formation, microtubule dynamics and chromosome segregation functions. All signatures have at least one substrate recognition component of E3 ubiquitin ligase complexes: *BTRC* in clusters 1 and 2; and *FBXW11* in cluster 3. Cluster 1 has genes involved in spindle checkpoint (*BUB1, CDC20*). The cluster 2 signature has many genes related to DNA repair (*CETN2, FANCB, H2AFX, ERCC1, ERCC4, PARP1, XPA*) and DNA replication (*RPA2, MCM10*). Moreover, it has three important genes in the signaling path that activates the *STAT3* transcription factor: *SRC, NFKB1* and *IL6R*. Indeed, the *STAT3* gene expression levels are significantly higher in cluster 2 (ANOVA p-value: 5.58 x 10^−15^) both in comparison with cluster 1 (T-Test p-value: 1.08 x 10^−9^) and cluster 3 (T-Test p-value: 1.14 x 10^−8^) patients (see Supplementary Figure 6). The cluster 3 signature contains genes involved in three different apoptotic mechanisms: induced by TNF-α (*TNFRSF1A* and *BAG4*), induced by Endoplasmatic Reticulum stress (*CAPN1* and *CAPN2*) and caspase-independent apoptosis (*ENDOG*).

**Table 4.** Common biological categories present in the gene signatures. All cluster signatures have genes that can be grouped in the following categories: NF-ĸB signaling, chromosomal instability and ubiquitin-proteasome system.

|  | **NF-κB Signaling** | **Chromosomal Instability** | **Ubiquitin-Proteasome System** |
| --- | --- | --- | --- |
| Cluster 1 | BTRC, CUL1, SRC, NFKBIA, NFKB1, NFKB2, REL, RELB, CHUK | CDC20, BUB1, MLFPIP, CENPC1, MIS12, PMF1, NDC80, RAD21, STAG1 | BTRC, CUL1, PSMB9, PSMC2, PSMF1 |
| Cluster 2 | BTRC, SRC, NFKBIA, TNFRS10B, IL6R | MIS12, DSN1, MLFPIP, CENPC1, PLK1, NEDD1, TUBGCP5, TUBGCP6 | BTRC, PSMB3, PSMC3 |
| Cluster 3 | FBXW11, AKT2, TNFR1A | CDC16, CDC27, NEDD1, TUBGCP5, KIF2B, KIF2C | FBXW11, PSMC3, PSMD9 |

Then, we searched for possible relationships between the gene signatures and genes commonly mutated in the studied tumours. We observed that some signature genes also present somatic mutations (*REL* and *RAD21* in cluster 1, *ERCC4* and *XPA* in cluster 2, and *AKT2* in cluster 3) or that mutated genes are direct neighbors of the signature genes in the network (see Figures 2, 3, 4). A permutation test over the signature labels (see Methods) reveals a significant proximity of signature genes to mutated genes for cluster 1 and cluster 2 (p-value=8.76 x 10^−4^ and p-value=6.9 x 10^−3^ respectively, Supplementary Figure 9). For the particular case of cluster 3, only one mutated gene is present in the network and it is successfully selected as a signature gene. These outcomes highlight the strict relationship between signature genes and key processes in tumour development (in analogy with the network-based approach of Novarino et. al.^13^).

Since the signature genes are the most central nodes in each cluster, we hypothesized that they might be suitable drug targets. For this purpose we collected, from the DrugBank database, the drugs that target genes in the signatures (Supplementary Table 1) and we evaluated in the ClinicalTrials repository if these drugs are under ongoing clinical trials for cancer treatment. We observed that 11 genes from the cluster signatures are being tested: 4 and 3 genes, from cluster 1 and 2, respectively; 3 genes from both cluster 1 and 2; and 1 gene from both cluster 1 and 3 (Table 5, Supplementary Table 2).

**Table 5.**
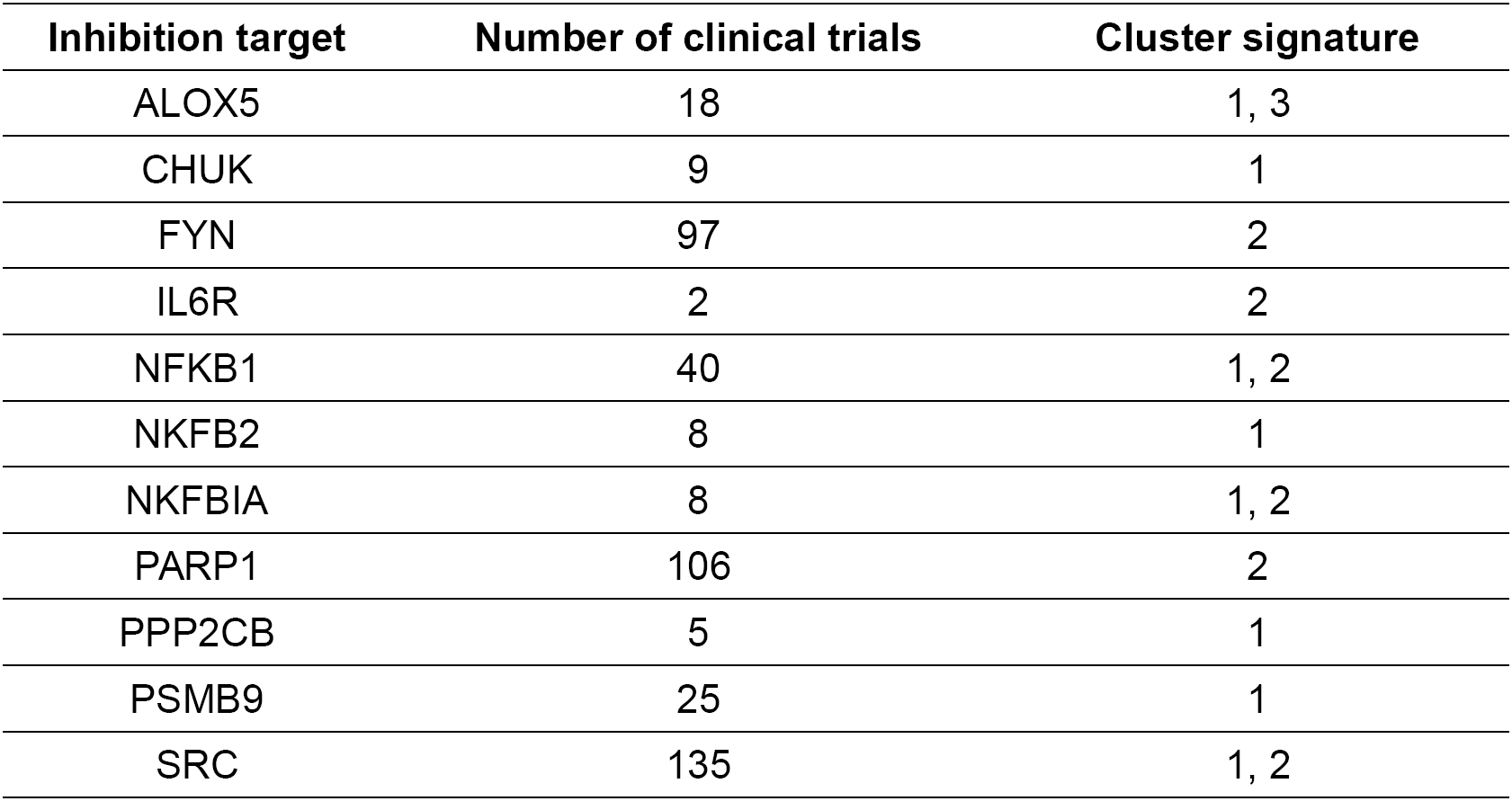
– List of genes from the signatures that are also being tested in ongoing clinical trials studies (according ClincalTrials.gov).

| Inhibition target | Number of clinical trials | Cluster signature |
| --- | --- | --- |
| ALOX5 | 18 | 1, 3 |
| CHUK | 9 | 1 |
| FYN | 97 | 2 |
| IL6R | 2 | 2 |
| NFKB1 | 40 | 1, 2 |
| NKFB2 | 8 | 1 |
| NKFBIA | 8 | 1, 2 |
| PARP1 | 106 | 2 |
| PPP2CB | 5 | 1 |
| PSMB9 | 25 | 1 |
| SRC | 135 | 1, 2 |

We then asked whether the expression level of the signature genes could predict the patients survival in each cluster, independently of the tumour type. For cluster 1 and 3, survival information were available only for 17 and 32 patients, respectively, which resulted in non-significantly different survival curves, possibly due to the low power of the test (see Supplementary Figures 7 and 8). For cluster 2, we retrieved the clinical information for 448 patients: the survival curves showed that the gene signature significantly separated the patients in two groups according to good or bad survival outcome (Log-rank test p-value = 4.54 x 10^−3^, see Methods and Figure 5).

**Figure 5.**
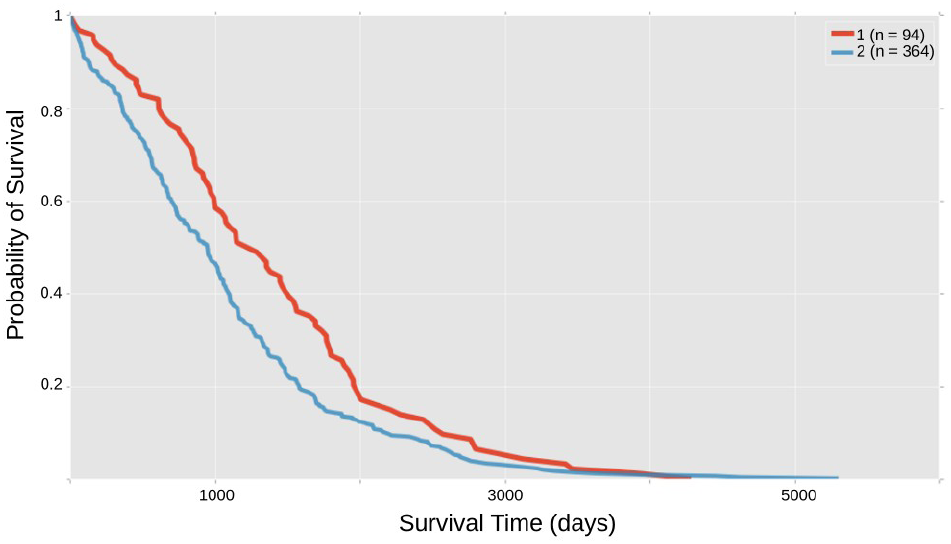
Gene signature and survival outcome. The cluster 2 signature defined two groups of patients with significantly different Kaplan-Meier survival curves.

We tried to translate our results into novel therapeutic strategies by applying, for a subset of tumours in cluster 2 (which contained the largest and most heterogeneous set of tumours), a set of drugs on targets taken from the signature and from related biological functions. We selected three drugs: Bortezomib, for targeting the proteasome and the NF-ĸB pathway; BI6727, for targeting the cluster 2 signature gene *PLK1;* and the PF-00477736 drug, to target the *CHK1/2* genes, which are not in the signature, but also plays a role in the DNA damage response. We tested these drugs, alone or in combination, on the glioblastoma cell line T98G and the breast adenocarcinoma model MCF-7. Both cell lines were highly sensitive to Bortezomib, with an IC50 of 200 nM for MCF-7 and 0.6 nM for T98G (Figure 6). BI6727 treatment reduced viability in a concentration-dependent manner in both models, with the glioblastoma model showing increased responsiveness (IC50 of 69.2 nM *versus* 1.8 μM for MCF-7, Figure 6). Moreover, both cell lines showed low response to *CHK1/2* inhibition, with IC50 of 26.9 μM for MCF-7 and 15.1 μM for T98G (Figure 6). We then asked whether these drugs might synergize in the selected models. Although the combinations of PF-00477736 with either BI6727 or Bortezomib did not show any additive or synergistic effect in both cell lines (data not shown), we observed a cooperation effect between inhibition of PLK1 and proteasome activity (Figure 7A-B). Indeed, we observed in the treatment with the drug combination that the cell viability was significantly lower compared with single agent treatments in MCF-7 cells (Figure 7A, p < 0.05), showing a general additive effect (Supplementary Table 4). We observed low Combination Index values (< 1) for both cell lines, indicating synergistic effect for all concentrations tested in the breast cancer model, and for selected concentrations in the glioblastoma model (Figure 7, Supplementary Tables 3 and 4).

**Figure 6.**
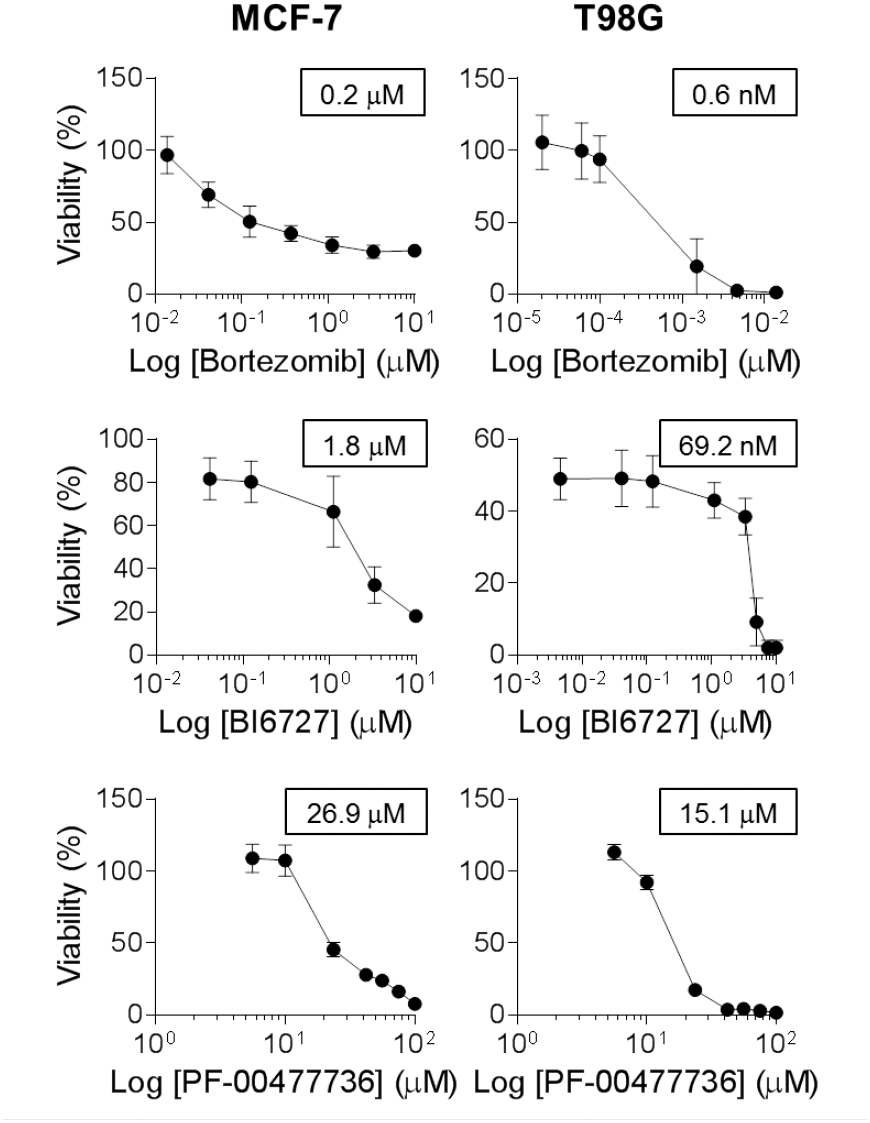
*In vitro* response of cancer cell lines from signature 2 to treatment with Bortezomib, BI6727 and PF-00477736 as single agents. MCF-7 and T98G cells were treated with increasing doses of Bortezomib (0.01 to 10 μΜ for MCF-7, 0.02 to 10 nM for T98G), BI6727 (0.04 to 10 μM for MCF-7, 0.004 to 10 μM for T98G), PF-00477736 (5.6 to 100 μM) and cell viability was measured 72h after drug administration by WST-1 assay (three independent experiments). Cell viability is represented as (mean ± SEM). IC50 values are reported in the boxes (GraphPad Prism 6).

**Figure 7.**
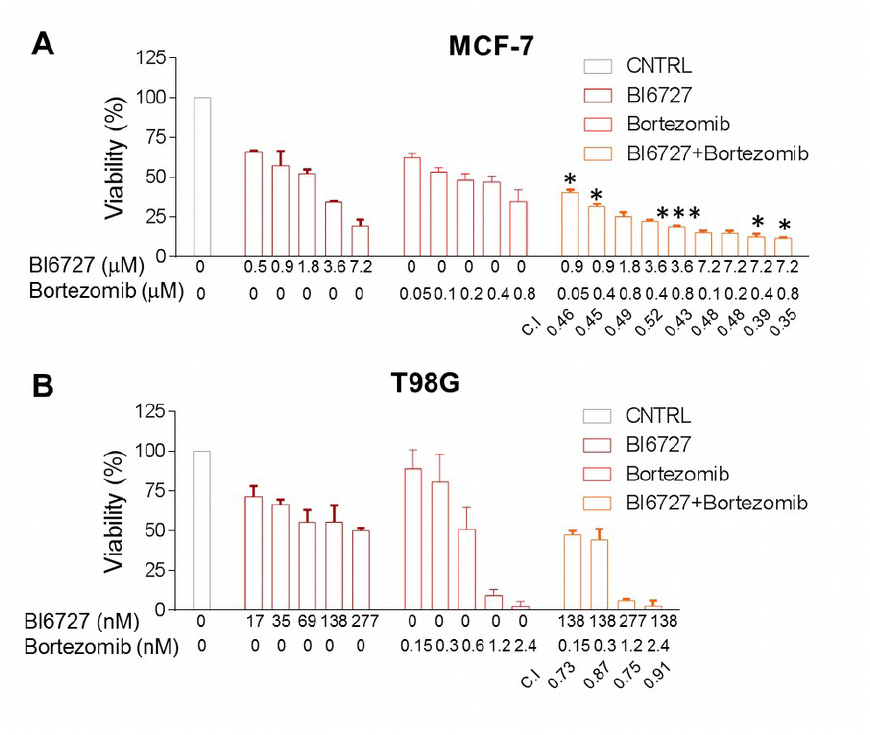
Sensitivity of MCF-7 and T98G cells to combined inhibition of PLK1 and proteasome activity. MCF-7 and T98G cells were treated with increasing doses of Bortezomib (0.05 to 0.8 μM for MCF-7, 0.15 to 2.4 nM for T98G) and BI6727 (0.5 to 7.2 μM for MCF-7, 17 to 277 nM for T98G), alone or in combination and cell viability was measured 72h after drug administration by WST-1 assay (three independent experiments). Statistical significance was determined by Student’s t test (*, P < 0.05; ***, P < 0.001). Combination index (C.I.) was calculated by CompuSyn software. (A) MCF-7 cells: combinations with a C.I. lower than 0.5 are shown. (B) T98G cells: combinations showing synergistic effect are shown.

## Discussion and conclusion

We studied the expression profiles of 11 tumours by considering a selected set of genes from the Ontocancro database and the BioPlex protein-protein interaction network. This knowledge-based selection reduced the dimensionality of the data to a highly curated list of cancer-related genes, involved in pathways that are hallmarks of cancer as cell cycle, inflammation, and apoptosis^14^. This approach also ensured that all studied genes had protein-protein interaction annotations, which are crucial to the understanding of how the signaling transduction propagates in the cell^15^. We clustered tumours by their gene-gene relationships defined by the Pearson’s correlation matrices, to evaluate the functional relationships between genes and their impact on transcriptome organization^16,17^. tumours from the same organ tended to group together, in agreement with previous studies showing that tissue-of-origin features provide the dominant signals in the identification of cancer subtypes^2,18^. However, the clustering also grouped tumours originating from different tissues, according to similarities in genomic alterations, as in the case of BRCA, OV, LUSC, and UCEC, which share common characteristics as presence of *TP53* mutations and multiple recurrent chromosomal gains and losses^3^. In particular, BRCA and UCEC grouped into a well defined subcluster, which may reflect their better prognosis when compared to other 10 tumour types^2^.

We integrated different types of biological information by a network approach, that allowed us to identify functional modules and to rank genes as network elements^19,20^. We created a network for each cluster (starting from a common template of protein interactions and superimposing cluster-specific correlation profiles) and obtained specific gene signatures based on node ranking by centrality measures. These signatures presented genes mainly involved in three biological processes: NF-ĸB signaling, chromosomal instability and the ubiquitin-proteasome system (Table 4). The NF-ĸB signaling pathway regulates genes that participate in cell proliferation, innate and adaptive immune responses, inflammation, cell migration, and apoptosis regulation processes. The aberrant activity of NF-ĸB may act as survival factor for transformed cells which would otherwise become senescent or apoptotic^21^. The genes classified into the chromosomal instability category involve kinetochore formation, microtubule dynamics and chromosome segregation functions. The dysfunction in these genes may cause cell inability to faithfully segregate chromosomes, generating genomic alterations as DNA mutation, chromosomal translocation, and gene amplification. The mutant genotypes may confer beneficial phenotypic traits to cancer cells, such as sustained proliferative signaling and resistance to cell death^14^. Two genes classified into this category have already been related to clinical practice: the prognostic marker KIF2C^22,23^; and the *BUB1* gene, which expression correlates with poor clinical diagnosis^24,25^. The ubiquitin-proteasome system is the major degradation machinery that controls the abundance of critical regulatory proteins. Perturbation of the regulatory proteins turnover disturbs the intricate balance of signaling pathways and the cellular homeostasis, contributing to the multi-step process of malignant transformation^26^. Proteasome inhibitors have become valuable tools in the treatment of certain types of cancer, mainly because malignant cells show greater sensitivity to the cytotoxic effects of proteasome inhibition than non-cancer cells^27^.

In addition to common features, cluster 2 signature has several genes related to DNA repair (*CETN2, FANCB, H2AFX, ERCC1, ERCC4, PARP1, XPA*) and DNA replication (*RPA2, MCM10*). Interestingly, the tumours in this cluster usually present high rates (50% to 90%) of samples with mutated *TP53*, which is an important sensor for the cell DNA damage response^2,4,28^. The cluster 2 signature also presents the genes *SRC, NFKB*, and *IL6R*, which participates in the activation of *STAT3*, a transcription factor that is necessary for cell transformation^29^. We observed that *STAT3* gene expression is higher in the tumours of cluster 2 when compared with the tumours of clusters 1 and 3 (Anova p-value: 5.58 x 10^−15^). The cluster 3 signature has genes involved in three apoptotic mechanisms, which are induced by TNF-α (*TNFRSF1A* and *BAG4*), or Endoplasmatic Reticulum stress (*CAPN1* and *CAPN2*) and caspase-independent apoptosis (*ENDOG*). As the regulation of cell death serves as a natural barrier to cancer development, these processes may reflect different strategies that these tumours use in response to various cellular stresses.

Since the transcriptional disturbances observed in cancer can sometimes be explained by underlying somatic mutations^30,31^ we retrieved TCGA mutational data, and focused on cancer related mutations reported in the Catalogue of Somatic Mutations in Cancer (COSMIC) database. Many signature genes resulted also somatically mutated, or first neighbours to mutated genes (Figures 2, 3, 4), showing their strict relationship and the functional relevance of the biologically processes they are involved in.

In addition, several genes in the signatures or in their direct network neighborhood are already under clinical investigation in a variety of tumour conditions (as annotated in Clinicaltrials.org database). For example, the AKT pathway has been described as a potential drug intervention in clear cell renal carcinoma^32^: *AKT2* gene belongs to the signature of cluster 3 (comprising LGG, KIRC, and KIRP), it is somatically mutated in the tumours of cluster 3 and it has been annotated as drug-target according to the Drug Bank database.

We also asked whether the gene signatures could predict survival outcomes in each cluster, thus independently on the tumour type. Our results show that in cluster 2 (the only one with enough available samples) the gene signature defined two groups of patients with significantly different Kaplan-Meier survival curves (log-rank test p-value: 4.54 x 10^−3^).

Finally, we tested 3 existing drugs (targeting 2 genes belonging to cluster 2 signature, and 1 involved in a related biological process, but not directly belonging to the signature) on 2 tumour types of the cluster, T98G and MCF-7 models. PF-00477736 drug (a *CHK1/2* inhibitor, not in the signature)^33^ had poor effect on both cell lines, but they resulted highly sensitive to BI6727 (an inhibitor of the signature gene PLK1^34^) and to Bortezomib (proteasome activity inhibitor^35,36^), with a significant synergic action at several dosages, suggesting a novel therapeutic strategy to be further explored in preclinical models of cluster 2 tumours.

These observations indicate that our study succeeded in: 1) clustering tumours highlighting common functional mechanisms related to their transcriptional profile, and 2) selecting genes with a relevant functional role in the studied tumours, thus amenable of drug targeting. The combination of these results may thus provide the rationale for choosing novel drug targets and drug combinations, or for repurposing existing drugs towards tumours of the same cluster. As a possible future direction, once obtained an enlarged list of novel and repurposed drugs, the specific transcriptional and mutational profile of single patients, prioritized onto our signatures, might suggest specific combinations of drugs for a more targeted and personalized therapeutic approach.

## Methods

### Gene expression datasets

The gene expression datasets used in this study were retrieved from The Cancer Genome Atlas (TCGA) Data Portal, and included Agilent expression arrays of 2378 samples from 11 tumour types, with a different number of samples each (from 16 to 595, see Table 1). We selected for our analysis the genes from the BioPlex protein-protein interaction network^11^ (n=10961) that were also present in the Ontocancro database (n=1104), resulting in a list of 760 cancer-associated genes related to specific biological functions (such as cell cycle control, DNA damage response, and inflammation).

### tumour clustering

For each tumour dataset, we calculated a correlation matrix containing pairwise Pearson r_¡j_ coefficients between genes across all samples available for the tumour. In order to eliminate false correlations and indirect influences, the absolute correlation values (|r_ij_|) were adjusted with the Context Likelihood of Relatedness (CLR) algorithm^37,38^ implemented in the R/Bioconductor package ‘minet’^39^. The matrices were clustered using hierarchical clustering analysis (with Ward linkage) based on the element-wise Euclidean distance between each pair of tumour matrices A and B, calculated as follows:

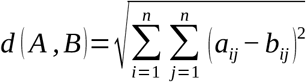

where *a_ij_* is the correlation between the genes *i* and *j* in the tumour A and *b_ij_* is the correlation between the genes *i* and *j* in the tumour B.

### Multi-tumour gene signatures

A network approach was applied to find gene signatures that characterized the clusters of tumours. First, we created a backbone network (BioPlex-Ontocancro) by selecting the genes present in the BioPlex protein-protein interaction network that were also annotated in the Ontocancro database. Then, for each cluster the gene-gene correlation coefficients r_ij_ were computed, and their absolute values |r_ij_| were adjusted with the CLR algorithm, producing the z_ij_ scores^37,38^. Each score matrix was superimposed to the BioPlex-Ontocancro network, producing three weighted networks (one for each cluster) in which genes were linked only if having correlated expression profiles (with weights given by positive z_ij_ scores, specific to each cluster) and a physical interaction at protein level (given by Bioplex-Ontocancro network, common to all clusters). We remark that the three cluster-related networks result different because of different weight values, or missing links (due to negative z scores set to zero). The networks were analyzed and visualized by Networkx Python package, Matlab and Cytoscape^40^.

For the networks of clusters 1 and 3, we selected the giant components (245 and 244 nodes, respectively), and for the cluster 2 we selected the two biggest components (149 and 118 nodes). After this selection, we retrieved a gene signature for each cluster composed by the most central genes (nodes), which were defined as those having the Spectral Centrality^12^ topological measure SC above the 90^th^ percentile. The SC calculates the effect of node removal on the network diffusivity based on the spectral properties of the Laplacian graph, and it has already been applied successfully to biological data such as the Immune System mediator network. Different results were obtained by considering Betweeness Centrality or weighted degree (Strength *W*) as centrality measures, as shown in Supplementary Table 5.

### Validation of the multi-tumour gene signatures

We evaluated the relevance of the genes in the signatures by several approaches.

First, we verified the proximity with the somatic mutational data extracted from the TCGA data portal for the considered tumours. To avoid cancer unrelated mutations, we considered only mutations that were reported also in the Catalogue of Somatic Mutations in Cancer (COSMIC) database^41^. We checked whether the signature genes had been reported as somatic mutated or if they occurred in the neighborhood of mutated genes in the networks. To quantify the proximity of gene signatures to mutated genes we located the nearest mutation (in terms of shortest paths on the network) for each signature gene, resulting in a collection of minimum distance values for each cluster. The average minimum distance from the mutated genes 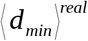was then calculated for each cluster and tested with a permutation test. We performed 10^6^ permutations of the signature labels and recalculated the average minimum distance of each new signature from the mutated genes. The p-values were calculated as

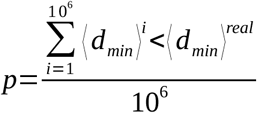

The results of the proximity analysis are reported in Supplementary Material Figure 9: the signatures of cluster 1 and cluster 2 are significantly closer to mutated genes than expected (p-values 9 x 10^−4^ and 6.9 x 10^−3^, respectively). The permutation test for cluster 3 is not completely meaningful because there is only one mutated gene, that anyway results to be one of the signature genes.

Secondly, we retrieved from the DrugBank^42^ (http://www.drugbank.ca/) and Drug Gene Interaction (DGIdb)^43^ databases which genes in the signatures were also mapped as drug targets. Third, we checked in the Aggregate Analysis of ClinicalTrials.gov (AACT) database (https://www.ctti-clinicaltrials.org/aact-database) for the existence of ongoing clinical trials evaluating the inhibition of genes in the signatures. Fourth, the prognostic potential of each gene signature was evaluated by considering the clinical data (days to death) available in the TCGA data portal. The patients having clinical information were clustered according to the expression levels of the gene signatures by using the k-means algorithm (Python package ‘scikit’), considering two patient groups: good versus bad survival outcome. Survivals curves were calculated for both groups: we applied the Kaplan-Meier method and evaluated their significance with the log-rank test (Python package “lifelines”). Fifth, we tested the effect of drugs inhibiting genes in our signatures or strictly related to them. The glioblastoma T98G and the breast adenocarcinoma MCF-7 cell lines were obtained from ATCC and DSMZ, respectively. Cells were cultured at a density of 10^5^ cells/ml in RPMI medium plus 10% FBS (plus 5% Sodium orthovanadate for T98G) for 72h with increasing concentrations of the following drugs: Bortezomib, BI6727, PF-00477736 (Selleckchem), alone or in combination. One hour and 30 minutes before the end of treatment, WST-1 reagent was added to the cell medium and cell viability was measured according to manufacturer’s instruction (Roche). The dose-effect response and the IC50 of each drug were calculated using GraphPad Prism 6 (GraphPad Software). To determine synergy, combination indexes were obtained with the CompuSyn software (ComboSyn Inc.): combination index values <1, =1, and >1 indicate synergism, additive effect and antagonism, respectively.

## Supplementary Information

Supplementary Figure 1 – Bioplex-Ontocancro Network. Network built from the 760 genes found both in BioPlex protein-protein interaction network and Ontocancro database

Supplementary Figure 2 – Overview of the cluster 1 network. Diamonds with red borders represent the genes in the cluster 1 signature and orange circles represent the mutated genes

Supplementary Figure 3 – Overview of the cluster 2 network. Diamonds with red borders represent the genes in the cluster 2 signature and orange circles represent the mutated genes

Supplementary Figure 4 – Overview of the cluster 3 network. Diamonds with red borders represent the genes in the cluster 3 signature and orange circles represent the mutated genes

Supplementary Figure 5 – Network of signature genes common to cluster 1 and 2. Even though the signatures can present common genes, they have different set of interactors in each cluster network.

Supplementary Figure 6 – Boxplot of STAT3 levels for clusters 1,2, 3. Cluster 2 patients presented higher STAT3 gene expression in comparison with cluster 1 (T-Test p-value: 1.08 x 10^−9^) and cluster 3 (T-Test p-value: 1.14 x 10^−8^).

Supplementary Figure 7 – Kaplan-Meier curves for the two groups of cluster 1 patients defined by K-means clustering approach. The clustering was applied only to the genes in cluster 1 signature. Only 17 patients had the survival information in the TCGA data portal (logrank-test pvalue = 0.9118)

Supplementary Figure 8 – Kaplan-Meier curves for the two groups of cluster 3 patients defined by K-means clustering approach. The clustering was applied considering only the genes in cluster 3 signature. Only 32 patients had the survival information in the TCGA data portal (logrank-test pvalue = 0.9056)

Supplementary Figure 9 – Plot of the distribution of the 10^6^ permutations for the 3 clusters (from left to right). The inboxes show the minimum average distances for the signatures (represented in the plots as red vertical lines), and the p-values with respect to the permutations.

Supplementary Table 1 – List of drug-gene interactions for the genes in the signatures, extracted from the Drug Gene Interaction database (DGIdb).

Supplementary Table 2 – List of ongoing clinical trials (according to ClinicalTrials.gov) that evaluate the inhibition of the genes in the signatures.

Supplementary Table 3 – Combination Indexes for BI6727 and Bortezomib treatment at different concentrations in the T98G cell line.

Supplementary Table 4 – Combination Indexes for BI6727 and Bortezomib treatment at different concentrations in the MCF-7 cell line.

Supplementary Table 5 – Spearman’s rank correlation values for the centrality measures (Spectral Centrality SC, Betweenness Centrality BC, strength W) on the nodes for the 3 clusters. The results refer to the whole node list (“All”) or only to the signatures, obtained as the top 10% of the ranked measures (“90th”). We remark the drop in correlation when considering only the gene signatures obtained by the different centrality measures.

## List of Abbreviations

TCGA: The Cancer Genome Atlas
SC: Spectral Centrality
COAD: Colon Adenocarcinoma
READ: Rectum Adenocarcinoma
LUAD: Lung Adenocarcinoma
LUSC: Lung Squamous Cell Carcinoma
GBM: Glioblastoma Multiforme
OV: Ovarian Serous Cystadenocarcinoma
BRCA: Breast Invasive Carcinoma
UCEC: Uterine Corpus Endometrial Carcinoma
LGG: Brain Lower Grade Glioma
KIRC: Kidney Renal Clear Cell Carcinoma
KIRP: Kidney Renal Papillary Cell Carcinoma

## Authors’ Contributions

IFV, GM and GS contributed equally to the paper. GS, SB, IZ and DFD performed the experiments and interpreted the results. IFV, GM and JCM collected and analyzed the data. GM, GC and DR designed the research, interpreted the results, wrote the paper.

## Acknowledgements

We thank the Science Without Borders project of CAPES foundation (Ministry of Education of Brazil – Brasília -DF, Brazil) for the doctoral scholarship (grant number: 10186-13-1) for IFV. This study was supported by the Interomics CNR Flagship Initiative; Mimomics EU FP7 Project n. 305280; NGS PTL EU FP7 Project n. 306242; the CNPq Project n. 402547/2012-8; the Associazione Italiana per la Ricerca sul Cancro AIRC (Investigator Grant to GM, n. 19226); the Programma di ricerca Regione- Universita 2010-2012 (L. Bolondi); and the Innovative Medicines Initiative (IMI) 2 project “HARMONY”, n 116026.

